# Insights into Therapeutic Discovery Through the Kelch Domain Structure of Keap1 at Ambient Temperature

**DOI:** 10.1101/2024.07.30.605796

**Authors:** Merve Yilmaz, Belgin Sever, Yigit Kutlu, Mehmet Gul, Ceren Okuducu, Serra Tavli, Masami Otsuka, Mikako Fujita, Turkan Haliloglu, Halilibrahim Ciftci, Hasan DeMirci

## Abstract

The *Kelch-like-ECH associated protein 1* (Keap1) is a part of the E3-ubiquitin ligase complex that binds to *Nuclear factor erythroid 2-related factor 2* (Nrf2) protein and facilitates its degradation by the eukaryotic 26S proteasome. The Kelch domain of Keap1 includes six repeated structural signature motifs, approximately 45–55 amino acid residues in length. Each Kelch repeat contains highly conserved residues and is known to form one blade of beta-propeller structure. Here, we report the dimeric Kelch domain of Keap1 determined at 3.0 Å resolution at the Turkish Light Source *‘Turkish DeLight*’ at ambient temperature. Our structure provides new structural dynamics information of the dimeric Keap1 Kelch domain at ambient temperature. It displays potential conformational changes of Keap1 residues, particularly at the Dimethyl fumarate (DMF) and Nrf2 binding regions, due to observed temperature shifts. Supported by the Gaussian Network Model (GNM) analysis, the dynamics of the Kelch domain revealed that the allosteric behavior of DMF binding residues is fully established in the ambient temperature structure. We also performed complementary molecular docking studies using our ambient temperature structure for numerous compounds acting as electrophilic irreversible indirect or non-covalent direct inhibitors of the Keap1/Nrf2 complex. Our data suggest that our previously reported novel compound, a hybrid of *L*-carnosine and *L*-histidyl hydrazide (CNN), revealed the most favorable scoring functions and prominent interactions with critical Keap1 residues. Collectively, our *in silico* and *in crystallo* results suggest a new potential lead compound for Keap1 inhibition. Additionally, understanding the dimeric form of the Keap1 Kelch domain and conformational changes around the DMF and Nrf2 binding sites at ambient temperature is critical for understanding Keap1-Nrf2 interaction dynamics.

## Introduction

A substrate adaptor protein involved in the Cullin-3-dependent E3 ubiquitin ligase complex is called *Kelch-like ECH-Associated Protein 1* (Keap1). Keap1 is composed of 625 amino acids, 27 of which are cysteine residues, and has a molecular weight of approximately 70 kDa. The DGR/Kelch domain of Keap1, which includes evolutionary conserved six Kelch repeats, mediates *Nuclear factor erythroid 2-related factor 2 (Nrf2)*-Keap1 interaction. The cryogenic structure of the human Kelch domain of Keap1 has been previously determined at 1.85 Å resolution by synchrotron X-ray crystallography. The Kelch domain is formed by assembling six repeats of bladed β-propeller structures. Kelch repeats have highly conserved glycine, tyrosine, and tryptophan residues, which have essential functional roles ***(Li et al., 2004)***. Nrf2 binds to Keap1 through the Kelch domain from two different binding motifs, ’DLG and ETGE,’ in the Neh2 domain. The DLG (Asp-Leu-Gly) motif is found in the N-terminal region of Nrf2 and interacts with the Kelch domain. The ETGE (Glu-Thr-Gly-Glu) motif is also found in the Neh2 degron and interacts with the Kelch domain ***(Tong et al., 2006)***.

The human Keap1 is a critical part of the Keap1-Nrf2-ARE (Antioxidant Response Elements) pathway that regulates the genes involved in detoxification, cytoprotective processes, and antioxidant defense to sustain cellular homeostasis ***(Canning et al., 2015)***. Keap1 is essential in controlling the ubiquitination-mediated degradation of Nrf2, which prevents Nrf2 from translocating into the nucleus regularly. However, in response to the presence of electrophilic substances or oxidative stress, some cysteine residues in Keap1 can perturb the Keap1-Nrf2 interaction. The dissociation of Keap1 and Nrf2 allows Nrf2 to translocate to the nucleus, bind to AREs in the DNA, and activate the transcription of genes involved in antioxidant defense, detoxification, and cellular protection ***(Baird et al., 2011)***.

Dimethyl fumarate (DMF) is a prodrug known to activate the Nrf2 pathway ***(Linker et al., 2011)***. In adults, it is indicated for the treatment of relapsing forms of multiple sclerosis (MS), including clinically isolated syndrome, relapsing-remitting disease, and active secondary progressive disease ***(Brennan et al., 2015)***. DMF binds to two distinct regions of Keap1; the Nrf2 binding region (Site 1) and near the second Kelch repeat (Site 2). DMF and Keap1 interaction from Site 1 is an intermediate interaction, and this site is the central region of interaction between Nrf2 and Keap1 ***(Unni et al., 2021)***. These two binding sites show the DMF capability of Nrf2 function activation with covalent and non-covalent interactions. This indicates that DMF is a potential therapeutic agent when dealing with oxidative stress in the cell. In addition, DMF analogs such as monoethyl fumarate (MEF) and fumarate (FUM) also exhibit the same binding modes as DMF. The interaction of DMF analogs with Keap1 might help Nrf2 activation and deal with oxidative stress in a similar manner ***(Unni et al., 2021)***.

In this study, we determined the dimeric Kelch domain of Keap1 at ambient temperature. We investigated the dimerization of the Kelch domain and identified the residues contributing to dimerization. We compared our structure with previously published structures for DMF and Nrf2 binding residues. Computationally, we performed GNM and GNM-based Transfer Entropy (GNM-TE) analysis to understand the dynamic behavior of the Kelch domain structure at cryogenic and ambient temperatures, especially the dimerization and DMF binding behaviors.

We also conducted molecular docking studies on our structure with our previously reported compound CNN (a hybrid of *L*-carnosine and *L*-histidyl hydrazide) ***(Noguchi et al., 2019)***, *L*-carnosine and fumarate derivative. Our findings, combined with existing knowledge of the structure and function of the Kelch domain, may help to drive future research for understanding Keap1-Nrf2 dynamics and the discovery of new drugs affecting the Keap1-Nrf2-ARE pathway.

## Materials and Methods

### Transformation and Expression

Kelch domain of Keap1 with the codon-optimized human sequence was cloned into a pet28a plasmid, including an N-terminal hexahistidine tag with a thrombin protease cut site. As cloning restriction enzyme cut sites, NdeI and BamHI were chosen, and the kanamycin resistance gene was used as a selection marker. The constructed plasmid was transformed into a competent *Escherichia coli* (E. coli) Rosetta2™ BL21 strain as described in ***Ertem et al., 2022***. Transformed bacterial cells were grown in 6 liters of regular Lysogeny Broth (LB) media containing 50 µg/mL kanamycin and 35 µL/mL chloramphenicol at 37 °C. When the OD_600_ value reached to 1.2, the protein expression was induced using β-D-1-thiogalactopyranoside (IPTG) at a final concentration of 0.4 mM for 18 h at 18 °C. Cell harvesting was performed using a Beckman Allegra 15 R desktop centrifuge at 4 °C at 3500 rpm for 45 mins. The cell pellets were stored at −80 °C until protein purification.

### Protein Purification

The cells were sonicated in a lysis buffer containing 500 mM NaCl, 20 mM Imidazole, 50 mM Tris-HCl (pH 8.5), 5% glycerol, and 0.1% Triton-X100. The cells were homogenized by dounce glass tissue homogenizer and lysed using a Branson W250 sonifier (Brookfield, CT, USA). The cell lysate was centrifuged at 4 °C at 35000 rpm for 1 hour with a Beckman OptimaTM L-80XP Ultracentrifuge equipped with a Ti45 rotor (Beckman, Brea, CA, USA). The pellet containing membranes and cell debris was discarded. The supernatant containing the soluble protein was filtered through a 0.2-micron hydrophilic membrane and loaded to a Ni-NTA column that was previously equilibrated with an equilibrium buffer containing 200 mM NaCl, 20 mM Tris-HCl (pH 8.5), 20 mM Imidazole. The column was washed with a wash buffer to discard unbound proteins from the column. Afterward, the target protein, Keap1, was eluted using an elution buffer containing 250 mM NaCl, 20 mM Tris-HCl (pH 8.0), and 250 mM Imidazole.

### Crystallization

The crystallization screening of Keap1 was performed using the sitting-drop microbatch under the oil method against ∼3000 commercially available sparse matrix crystallization screening conditions in a 1:1 volumetric ratio in 72-Terasaki plates (Greiner Bio-One, Kremsmünster, Austria). The mixtures were covered with 16.6 µL 100% paraffin oil (Tekkim Kimya, Istanbul, Türkiye). The Terasaki plates were incubated at 4 °C and checked frequently under a stereo light microscope. The best Keap1 crystals were obtained within one month in condition Salt Rx-II #44 which contain 4.0M Ammonium Acetate, 100mM Sodium Acetate Trihydrate (pH 4.6) and Salt Rx-II #45 4.0M Ammonium Acetate, 100mM BIS TRIS Propane (pH 7.0) and Salt Rx-II #46 which contain 4.0 M Ammonium Acetate, 100mM Tris-HCl (pH 8.5) (Hampton Research, USA).

### Ambient Temperature Data Collection and Processing

Ambient temperature X-ray crystallographic data was collected using Rigaku’s XtaLAB Synergy R Flow XRD system, as described in ***Gul et al., 2022***. Multiple crystals were screened using the modified adapter of the *XtalCheck-S* plate reader. After selecting well-diffracting crystals, diffraction data were collected from two crystals. The duration of exposure time was optimized to minimize the potential radiation damage caused by X-rays. A total of 2 crystals were used to collect a complete diffraction dataset. Diffraction data were continuously collected for approximately 3 hours at 90 percent attenuation. The detector distance was set at 120.0 mm, while the scan width was 0.5 degrees, and the exposure time was 5.0 seconds per image. The diffraction data were set up in *CrysAlisPro* to complete the automated data collection. The collected data was then merged using the profit merge process with *CrysAlisPro* 1.171.42.59a software ***(Rigaku OxfordDiffraction, 2022***) to produce an integrated reflection dataset (*.mtz) file for further analysis, as described in ***Gul et al., 2022***.

### Structure Determination and Refinement

The structure was determined using molecular replacement with the *PHASER-MR* program implemented in the *PHENIX* suite, using the cryogenic synchrotron Keap1 crystal structure (PDB ID: 6ROG) as the initial search model **(*Adams et al., 2010*)**. Following rigid body and simulated annealing refinement, individual coordinates and TLS parameters were refined. Potential positions of altered side chains and water molecules were examined, and the models were manually constructed/reconstructed using the *COOT* program ***(Emsley et al., 2004)***. The final structure refinement was performed using *phenix.refine* in *PHENIX*. Structural figures were generated using *PyMOL*.

### Gaussian Network Model (GNM) Analysis

To analyze the structural dynamics of the Keap1 Kelch domain structures***, w***e employed the Gaussian Network Model (GNM) ***(Bahar et al., 1997; Haliloglu et al., 1997)***. GNM decomposes the fluctuations of residues in a protein structure into a series of orthogonal modes of motion. These modes range from highly collective global motions to localized fluctuations across the protein structure. The study focuses on the slow mode spectrum, particularly the 10 and 5 slowest modes for the dimeric and monomeric structures, respectively. These normal modes generally account for a significant portion of a protein’s functional dynamics.

The equilibrium correlation between the fluctuations of two residues i and j, respectively, ΔR_i_ and ΔR_j_, is given as:

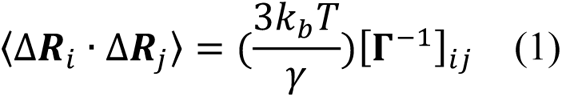

where **Γ** is a symmetric matrix of the Kirchhoff (connectivity) matrix with a r-cut of 10 Å to assume interactions, *γ* is the force constant of the Hookean pairwise potential function, representing the interactions between the residues in the folded structure. *T* is the absolute temperature in Kelvin, and *k_b_* is the Boltzmann constant. Eq. 1 can be rewritten as:

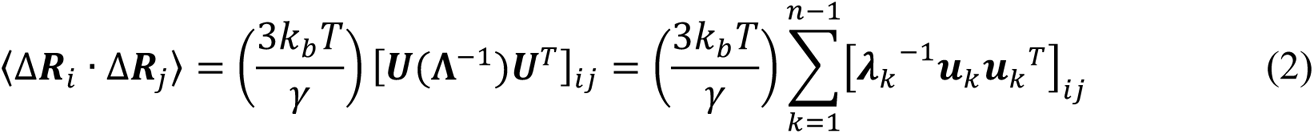

where *k* is the *k*-th vibrational mode in the spectrum of *n*-1 modes, *n* being the number of residues. ***U*** is an orthogonal matrix whose columns ***u****_i_* are the eigenvectors of **Γ**, and **Λ** is the diagonal matrix of the eigenvalues *λ_k_*.

Additionally, GNM-based Transfer Entropy predictions explore protein dynamics in causal interrelation.

GNM-TE combines Transfer Entropy (TE) with GNM to assess the directional flow of information between residues i and j ***(Hacisuleyman & Erman, 2017; Altintel et al., 2022; Ersoy et al., 2023)***. TE quantifies the reduction in uncertainty about the movements of residue j given the movements of residue i, incorporating a time delay τ, represented as T_i→j_(τ). Net TE values equate to the difference between T_i-j_ (τ) and Tj-i (τ)

The TECol score for each residue, referred to as residue i, is determined by multiplying its cumulative positive net TE value (the total of positive net TE values that residue i transmits to other residues) by its collectivity value K_i,s_ within each subset s.

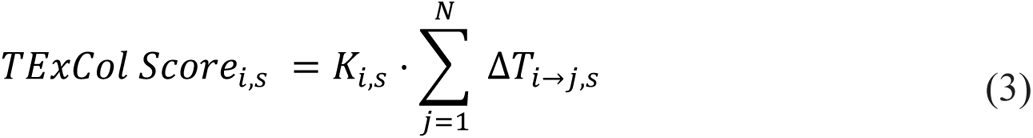

The collectivity value K_i,s_ within each subset s is calculated as

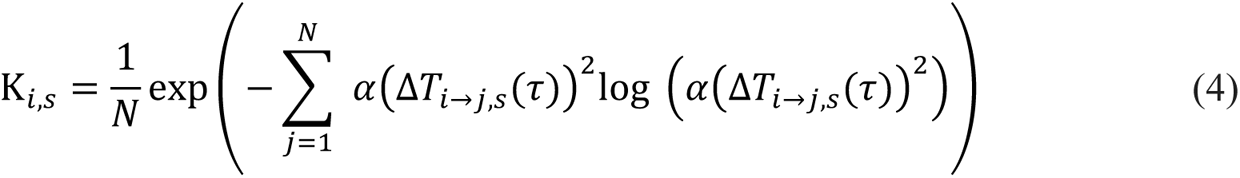

where s represents a selected subset of slow GNM modes, N is the total number of residues, and α denotes the normalization factor. The TECol score, combining both TE and K values, is essential for identifying the most functionally significant global information source residues. These residues have substantial influence and act as powerful effectors within the protein structure.

### P-value analysis

To demonstrate the statistical significance between functional residues (e.g., DMF binding residues) and those identified through TECol score analysis, we conducted a p-value analysis. We randomly selected the same number of residues as those identified by the TECol scores and calculated the overlap with the DMF binding residues. A DMF binding residue was considered a match if it was within 4 Å of any randomly selected residue. This process was iterated 1,000 times to generate a randomly distributed population and a cumulative distribution function (CDF) for the overlaps. The p-value was then calculated using the CDF for the overlap between the DMF binding residues and those identified by the TECol scores.

### Molecular Docking Studies

ZINC database (https://zinc.docking.org/) was used to obtain the proper structures for exploration of potential Keap1/Nrf2 inhibitors (fumarates and carnosic acid derivatives) and potential Nrf2 activators (*L*-carnosine and *L*-carnosine derivatives).

The crystal structure of the ambient temperature Keap1 Kelch domain, *Keap1^Ambient_APO^*, was acquired from the RCSB database *(ref:* https://www.rcsb.org/structure/8X34). The Protein Preparation Wizard was utilized to prepare the raw protein to be used in molecular docking. The missing chains were added automatically by *Prime* and the protonation state was calculated by *PropKa* at physiological pH. Afterward, the top-ranked potential receptor binding sites were identified using *SiteMap*. The docking grid was determined by grid generation picking the hit binding site compromising the specified residues (Asn382, Arg483, Gln530, Gly371, Phe577, Ser363, Tyr334, 525, 572, and Val369). The generated grid was used for further docking experiments.

On the other hand, compounds were sketched and cleaned in the *Maestro* workspace. They were prepared with energy minimization using the *OPLS 2005* force field at physiological pH by the *LigPrep* module. Then, the best-minimized structures were submitted to the docking experiments without further modifications. The flexible ligand alignment tool was applied to superimpose our structurally similar ligands. The self-docking experiment was performed to validate the docking protocol with DMF. The optimum structure (lowest energy) was performed for the self-docking procedure. After the obtained ligand was submitted to *Glide/SP* docking protocols, the same docking procedures were carried out for all designed compounds ***(Schrödinger Release 2016-2: Schrödinger, LLC: New York, USA, Ciftci et al., 2022; Guven et al., 2023)***.

## Results

### The Kelch domain of Keap1 is determined in dimeric form

Here, we determined that the 62.6 kDa dimeric Kelch domain of Keap1 consists of 572 amino acids. The Keap1 crystal belongs to the *P*2_1_2_1_2_1_ orthorhombic space group with α= 90, β= 90, γ= 90, a = 76.61, b = 77.76, and c = 213.29. The dimer structure of Kelch domain Keap1 was determined to be 3 Å resolution at ambient temperature at the Turkish light source ‘*Turkish Delight*’ ***(Atalay et al., 2023)***. The determined dimeric Kelch domain of the Keap1 structure was deposited to the Protein Data Bank with the PDB ID:8X34 (Keap1^Ambient_APO^).

Our study suggests that the structure of Keap1 has dimerized at the Kelch domain (**Figure 1A**). As a result of monomer alignment, there is a slight difference between chain A and chain B, with an RMSD score of 0.21 Å (**Figure 1B**). The residues that mediate the dimerization are shown in **Figure 1C and 1D**. A comparison of the dimeric Keap1^Ambient_APO^ structure at 3 Å and the dimeric Keap1^Cryo_APO^ (PDB ID: 6ROG) structure at 2.1 Å shows a difference between the two structures with an RMSD score of 0.92 Å. To confirm that this difference results from temperature shift, the Keap1^Cryo_APO^ structure was re-refined to the resolution range of the Keap1^Ambient_APO^ (3 Å). The re-refined structure was then realigned with the PDB deposited structure. From this perspective, the resolution difference was eliminated, which suggests that the variation may be caused by the temperature shift. Additionally, the pairwise alignment demonstrated the distance between residues based on alpha carbons in Keap1^Ambient_APO^ and Keap1^Cryo_APO^ **(Supplementary Figure 1)**.

**Figure 1.**
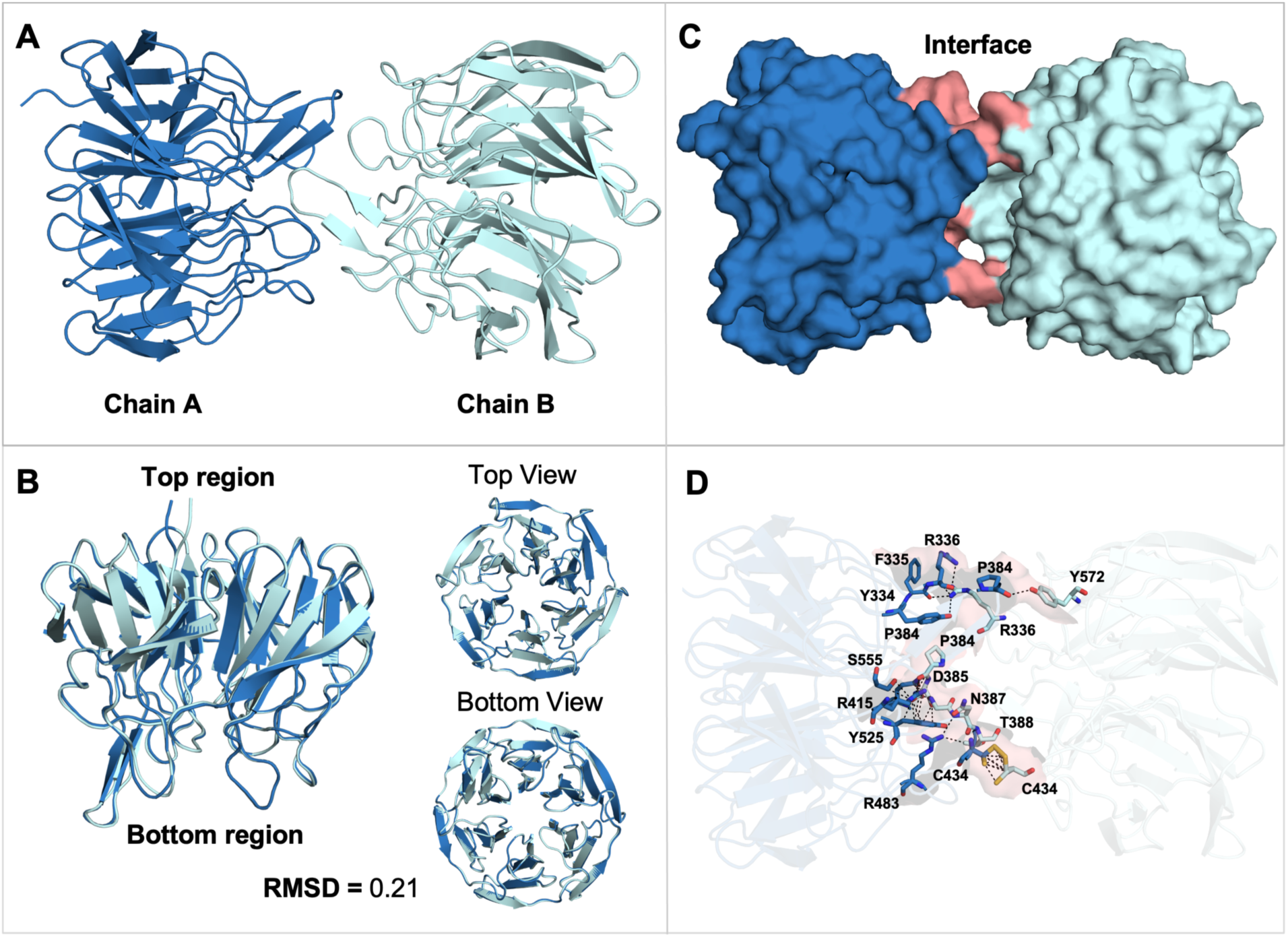
Dimerization of Kelch Domain and Alignment of monomers (A) Keap1^Ambient_APO^ dimerizes through its Kelch domain, with Chain A colored in sky blue and Chain B colored in pale cyan. (B) Alignment of Chain A and Chain B, with an RMSD score of 0.21 Å. The top and bottom views of the Keap1 are shown. (C) The protein interface between Chain A and Chain B is shown and colored in salmon. (D) Residues contributing to the protein interface and dimerization are highlighted and shown in sticks representation.

### Temperature shift causes conformational variations in DMF and NRF2 binding residues

Our structural analysis comparing the Keap1^Ambient_APO^ and cryogenic Keap1 structure in complex with DMF (Keap1^Cryo_DMF^) (PDB ID:6LRZ) structures revealed minor changes in the DMF interacting residues. These residues are found in the first Kelch repeat (Tyr334, Ser363, Val369, and Gly371), second Kelch repeat (Asn362), fourth Kelch repeat (Arg483), fifth Kelch repeat (Try525 and Glu530) and sixth Kelch repeat (Phe577 and Try572). We observed alterations on the top side Keap1 residues Asp382, Tyr334, Phe577, Tyr572, Glu530, Tyr525 and Arg483, and at the bottom side residues Val369, Gly371 (**Figure 2A**). Additionally, comparing both Keap1^Ambient_APO^ and Keap1^Cryo_APO^ demonstrates similar conformational changes with the Keap1^Cryo_DMF^ structure (**Figure 2B**).

**Figure 2.**
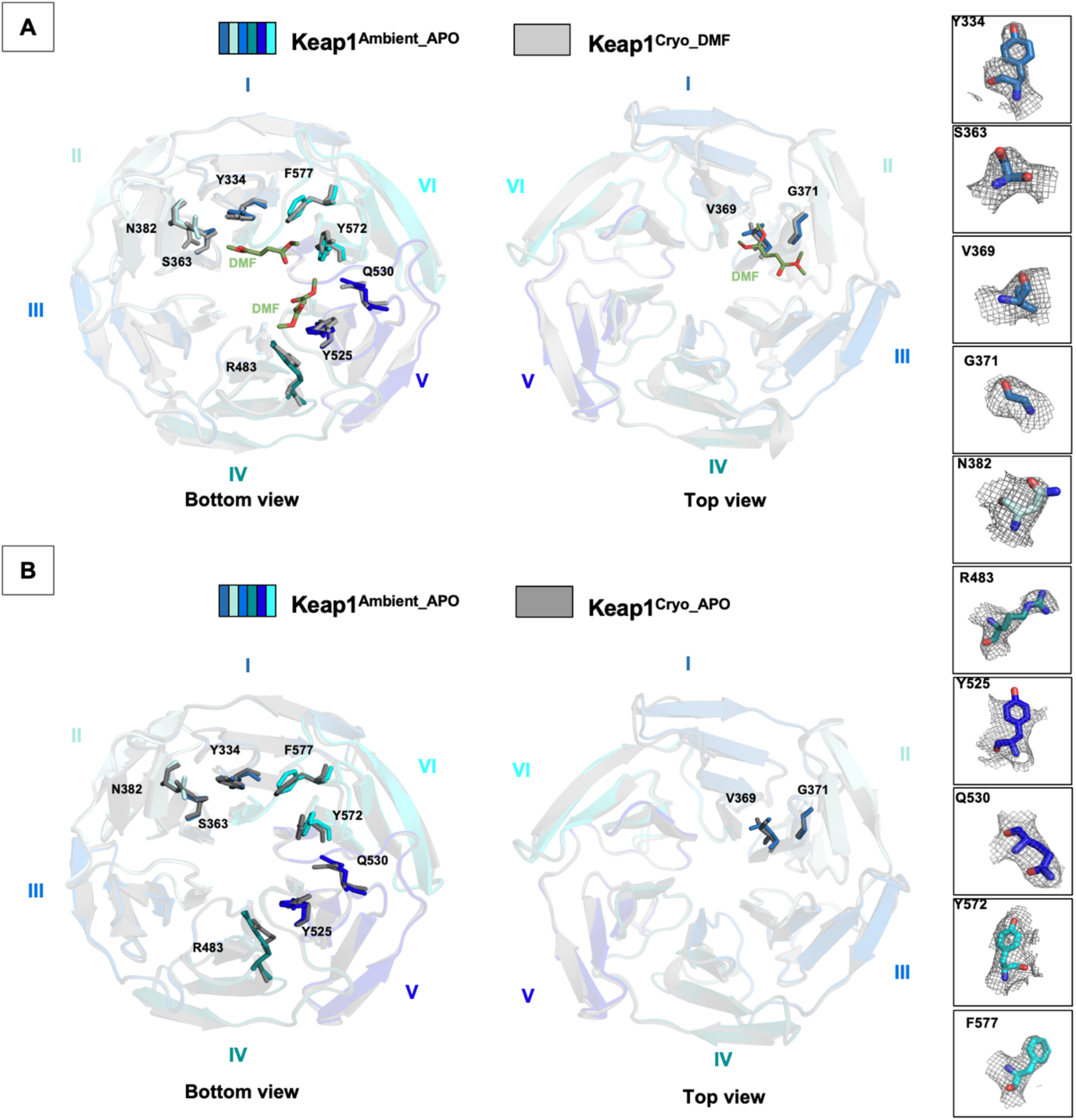
Investigation of residues interacting with DMF molecule in cryogenic and ambient structures (A) Superposition of Keap1^Ambient_APO^ and Keap1^Cryo_DMF^ (colored in gray80) is shown both from bottom and top views, with DMF binding residues displayed in sticks representation. (B) Keap1^Ambient_APO^ and Keap1^Cryo_APO^ (colored in gray50) are superposed, with DMF interaction residues also shown in sticks representation. DMF interaction residues are displayed with their respective electron density.

We compared our Keap1^Ambient_APO^ structure with cryogenic Keap1 structure in complex with ETGE peptide (Keap1^Cryo_ETGE^) (PDB ID: 5WFN), and cryogenic Keap1 structure in complex with ETGE peptide (Keap1^Cryo_DLG^) (PDB ID: 3WN7), to determine the conformational changes in Nrf2 protein interacting residues resulting from temperature shift. Our observation suggests major alterations in the residues Arg483, Ser508, Ser555, Arg415, Ser602, Ser363, Arg380, and Asn382, which interact with ETGE and DLG peptide motifs in the ambient structure (**Figure 3**).

**Figure 3.**
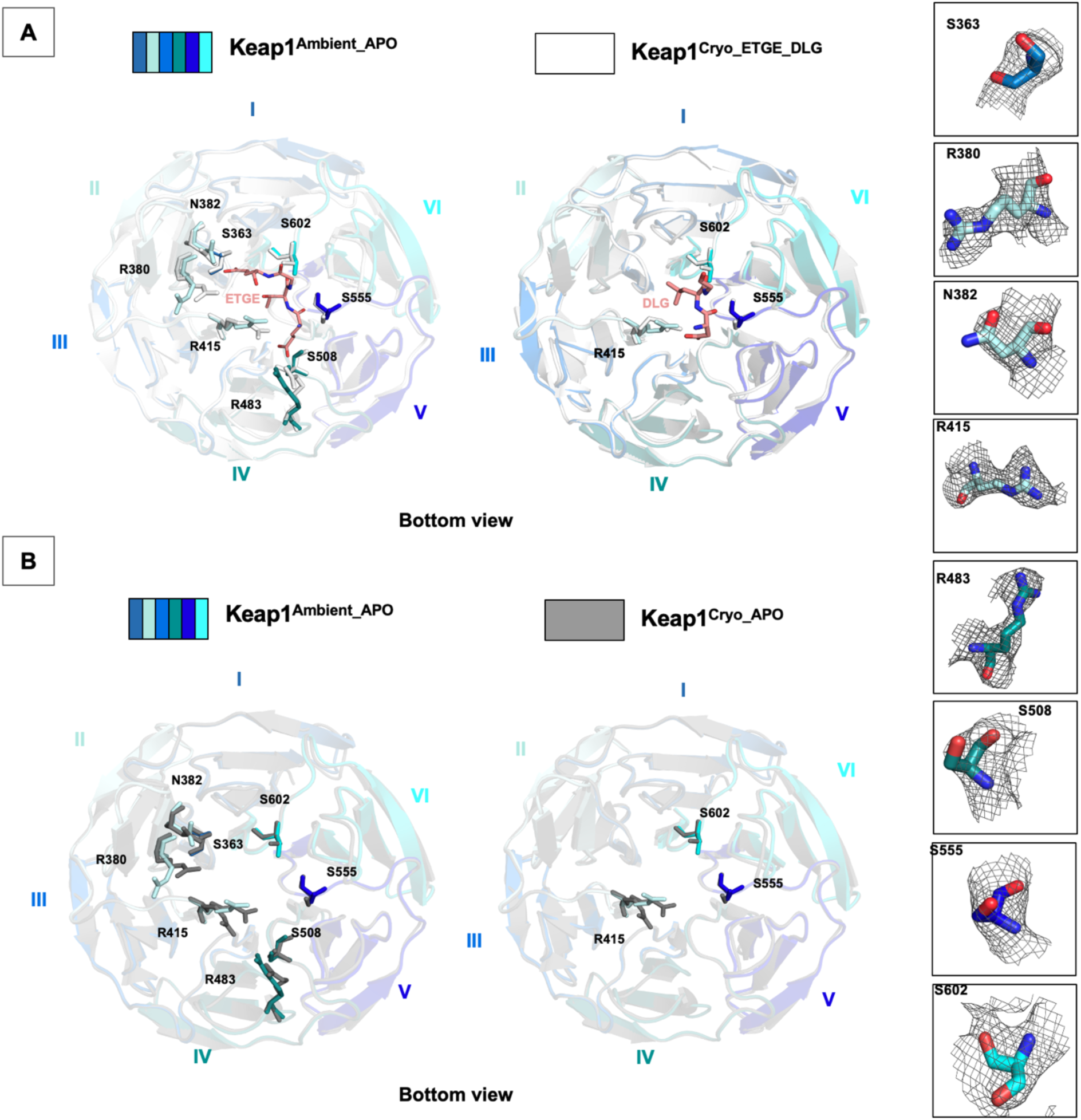
Residues interacting with NRF2 in Keap1^Cryo_APO^ and Keap1^Ambient_APO^ structures (A) Superposition of Keap1^Ambient_APO^ and Keap1^Cryo_ETGE^ (colored in white), with interacting residues displayed in stick representation. Superposition of Keap1^Ambient_APO^ and Keap1^Cryo_DLG^ (colored in white), with interacting residues displayed in stick representation. (B) Superposition of Keap1^Ambient_APO^ and Keap1^Cryo_APO^ (colored in gray50), with ETGE and DLG interacting residues also shown in stick representation. NRF2 interacting residues are displayed with their respective electron density.

### GNM analysis reveals differences in residue correlations between dimeric Keap1 structures

The structural dynamics of the Keap1 Kelch domain were investigated by examining residue cross-correlations and their differences for the ten slowest GNM modes of the Keap1^Cryo_APO^ and Keap1^Ambient_APO^ (**Figure 4**). The residue correlations were generally similar for both structures, with the residues predominantly correlating within their respective chains. However, residues Pro384, Asp385, and Gly386 involved in the dimerization (**Figure 1**) show correlated fluctuations with the other chain (**Figure 4A**). Additionally, we identified some variations in the correlations in specific regions, particularly near residues involved in dimerization (**Figure 1**), which are Arg336, Pro384, Asp385, and Gly386 for chain A, and Arg336, Pro384, Asp385, Gly386, and His575 for chain B (**Figure 5B**). However, Arg336 and Asp385 are observed to be the residues where the β-factor between two structures is the highest, suggesting that these differences possibly occur due to thermal fluctuations **(Supplementary Figure 2)**.

**Figure 4.**
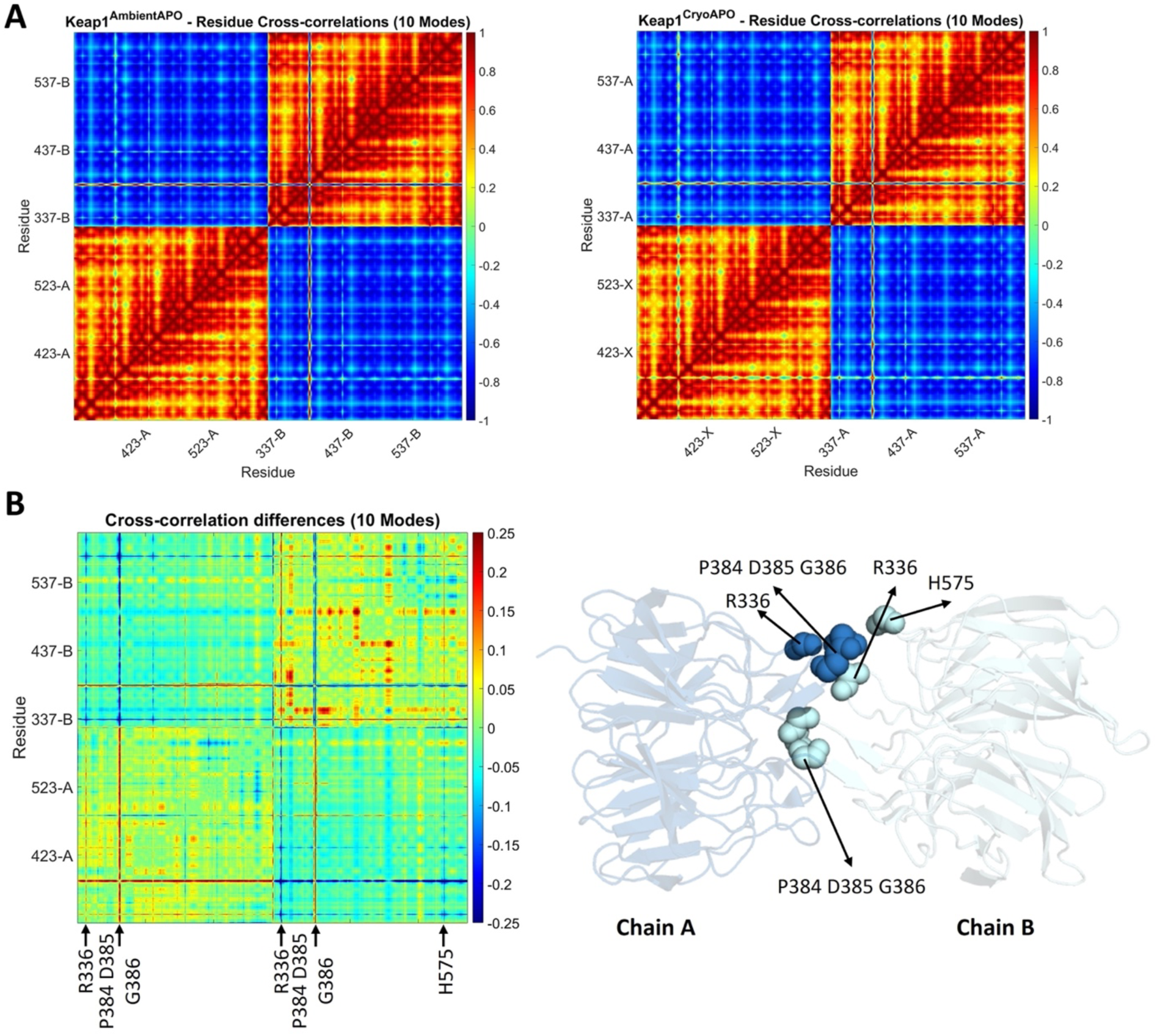
Cross-correlation maps for Keap1 apo structure in ambient and cryogenic conditions (A) GNM residue cross-correlations for ten slowest GNM modes of the Keap1^Ambient_APO^ and at Keap1^Cryo_APO^ (B) Cross-correlation differences of these structures. Residues that show differences in correlations are marked and displayed in a 3D structure where chain A is colored in skyblue and chain B is colored in pale cyan.

**Figure 5.**
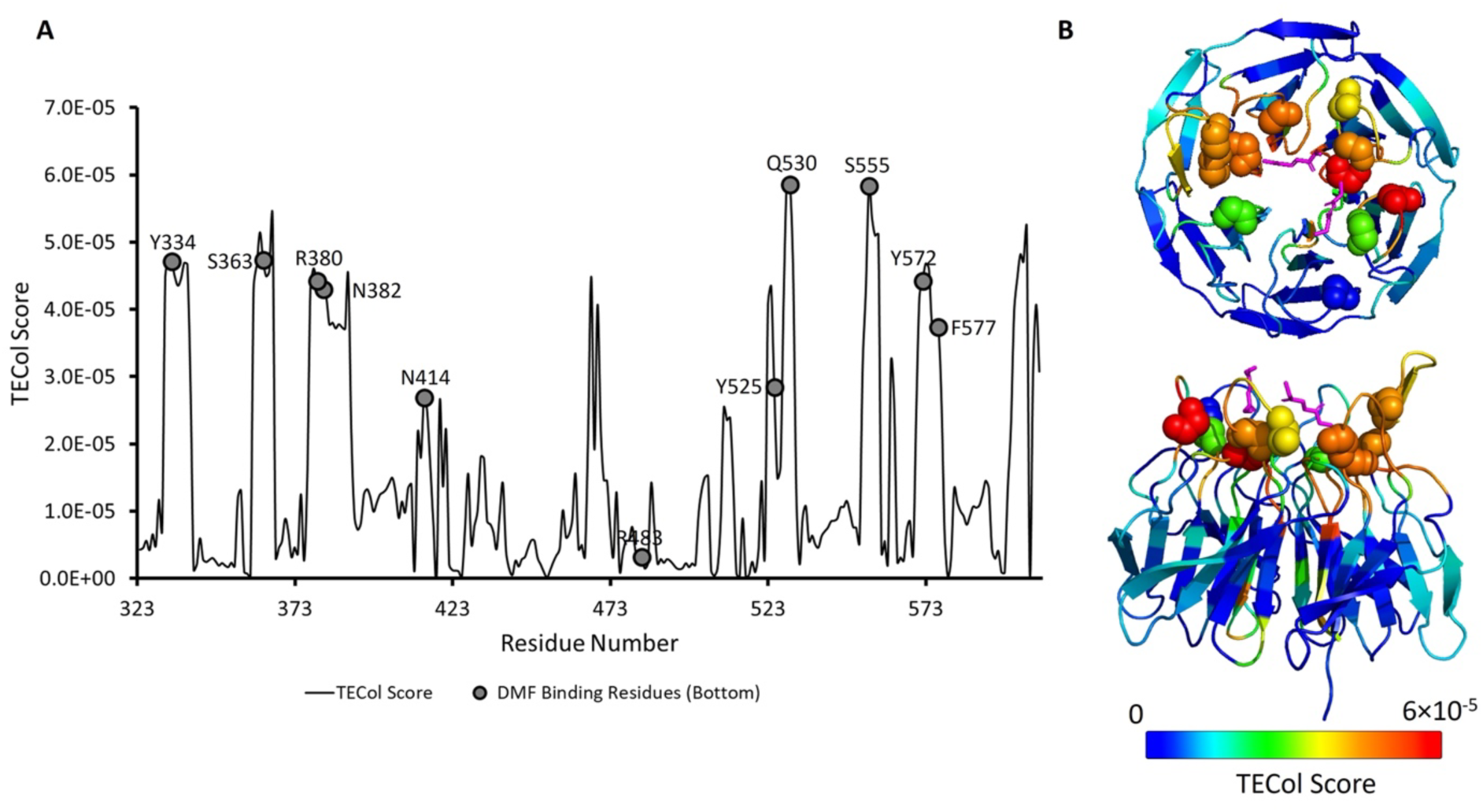
The TECol score results with the subset of slow modes consisting of GNM modes 3 to 5 for the monomer Keap1^Ambient_APO^ chain A. (A) Graphical representation of the TECol score results where bottom-side DMF binding residues are marked on the TECol score graph. (B) 3D representation of the TECol score results from two angles. Residues are colored according to the TECol score with respect to the rainbow spectrum. DMF binding residues are shown as spheres and DMFs are shown as magenta sticks.

In addition, similar analyses were conducted on the monomer Keap1^Cryo_DMF^ and Keap1^Ambient_APO^ structures. The correlation profiles for these structures were very similar, and no significant differences were observed in the correlation variations **(Supplementary Figure 3)**.

### Ambient temperature maximizes the allosteric capacity of the DMF binding residues in the Keap1 Kelch domain

In the previous sections, it was mentioned that temperature changes cause some conformational changes in the Keap1 Kelch domain. Here, these conformational changes of the monomeric Keap1 Kelch domains were converted into a difference vector, and then the magnitude between the C-alpha atoms of each residue after superposing the monomer structures Keap1^Cryo_DMF^ and Keap1^Ambient_APO^ (chain A). Subsequently, we investigated whether a GNM mode was similar to the profile of the magnitudes of residue difference vectors. Our calculations revealed that this profile highly correlated with the third slowest GNM mode (**Supplementary Figure 4)**. This means that this specific dynamic mode mainly induced the conformational fluctuations from the cryogenic to ambient temperature. Although the third mode of the two structures is similar **(Supplementary Figure 5)**, this mode may be considered significant for understanding functional dynamics.

Next, the GNM-TE method was applied to the monomeric structures Keap1^Cryo_DMF^ and Keap1^Ambient_APO^ (chain A), focusing on the five slowest GNM modes that represent the slow end of the dynamic spectrum. GNM-TE predicts the residues with the dynamic capacity to transfer information to other residues. These residues with high TECol scores are information source residues that collectively influence others and act as powerful effectors. In the five slowest GNM modes, we have used the subsets of 1 to 5, 2 to 5, and 3 to 5 slow modes. The subsets of slow modes 1 to 5 and 2 to 5 highlight the binding site residues to the molecule PG5 (see **Supplementary Figure 6** for GNM-TE results of a subset of modes 1-5 and 2-5.). However, in the subset of 3 to 5 slow modes, the DMF binding regions emerged as prominent effector residues (**Figure 5**).

Interestingly, upon the removal of the slowest and second slowest modes, the behavior of the third slow mode, jointly with the fourth and fifth slow modes, reveals the collective behavior of the DMF binding sites, which is otherwise latent in the subset of slow modes including the two global modes. This supports the functional significance of the third slowest GNM mode we observe upon the conformational fluctuations from the cryogenic to ambient temperatures. The TECol score successfully captures DMF binding residues at or near the peak positions (< 4 Å). Statistical significance analysis reveals a p-value of 0.002.

Additionally, when the same analysis was performed on the cryogenic structure, the results did not reveal the full dynamic capacity of the DMF binding sites **(Supplementary Figure 7)**. Although there are very slight differences between these modes in the two structures **(Supplementary Figure 5)**, it was observed that these small differences could result in variations in capturing functional movements of specifically the DMF binding regions. Furthermore, in the GNM-TE analysis for the ambient temperature structure, in addition to the DMF binding residues, Ser390, Val467, Asn469, and Gly605 also emerged as prominent effector residues. These findings imply that these residues are of allosteric importance, particularly for DMF binding behavior. The information flow graphs were also examined for residues Asn382 and Tyr334, critical for DMF binding (mentioned in the previous sections). It was observed that the information flow from these residues is very similar, and residues Tyr342, Val411, Tyr426, Ile506, Arg507, and Val514 are the ones receiving the most information transfer. These residues may also have an allosteric importance in the collective behavior induced by the DMF binding residues **(Supplementary Figure 8**).

### Molecular docking studies of fumarate and carnosic acid derivatives exhibit distinct binding profiles

The molecular docking analysis of 100 fumarate derivatives in the binding site of our structure revealed that 9 derivatives (ZINC IDs: 12433145, 14496485, 3860363, 5573987, 2383091291, 21945150, 5239687, 2265312934, 1911888877) were found to possess more significant affinity profile and docking scores compared to DMF and well-known fumarates such as itaconate (ITA), monomethyl fumarate (MMF), monoethyl fumarate (MEF), and fumarate (FUM). The docking score of the most effective derivative, ZINC 12433145, was detected as -9.130 kcal/mol compared to values of DMF, ITA, MMF, MEF, and FUM, which were found as -8.423, -7.533, -7.456, - 7.146, and -6.934 kcal/mol, respectively. ZINC 12433145 displayed crucial hydrogen bonding with key residues Asn382 and Tyr334 in a similar pattern with DMF in the binding site of the Keap1 Kelch domain (**Figure 6B**).

**Figure 6.**
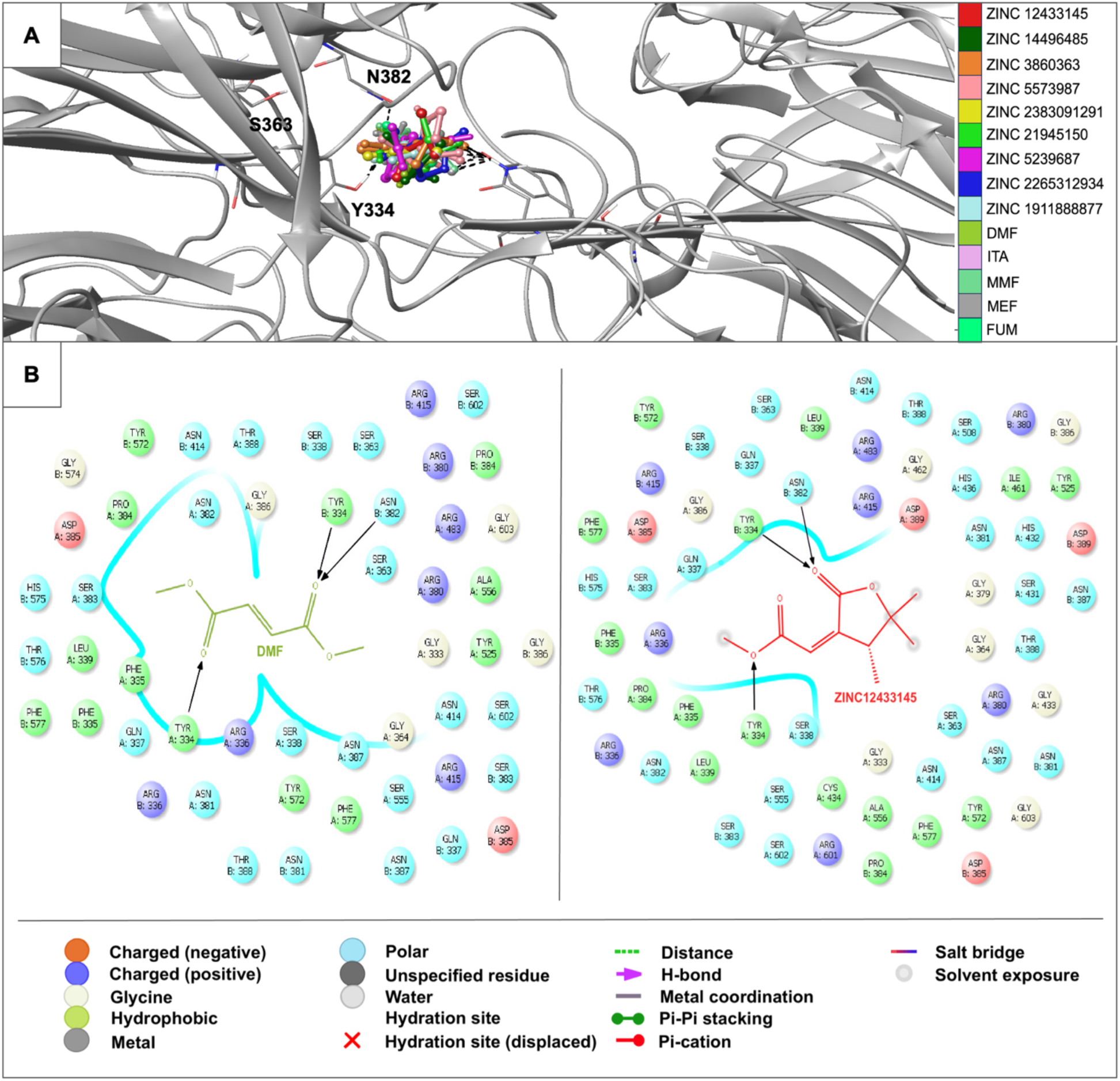
Docking results of fumarate derivatives and DMF, ITA, MMF, MEF and FUM in the binding site of Keap1^Ambient_APO^ structure. (A) Docking poses of fumarate derivatives and DMF, ITA, MMF, MEF and FUM in the binding site of Keap1^Ambient_APO^ structure. The light gray ribbon diagram in a metallic preset was displayed for all residues. The key residues were represented as sticks and colored by atom type: (C gray, O red, N blue). H-bonding was outlined with dotted black lines. (B) Docking interactions of ZINC 12433145 and DMF in the binding site of Keap1^Ambient_APO^ structure.

The carnosic acid and 100 carnosic acid derivatives exhibited a low binding profile compared with fumarate derivatives in the binding site of the Keap1 Kelch domain. Carnosic acid was identified as the most significant, with a docking score of -5.446 kcal/mol compared to the most promising carnosic acid derivative, ZINC 105508677 (-4.758 kcal/mol). Carnosic acid and ZINC 105508677 formed hydrogen bonding and π stacking with mainly arginine residues, indicating a distinct binding profile than fumarates (**Figure 7A and 7B**).

**Figure 7.**
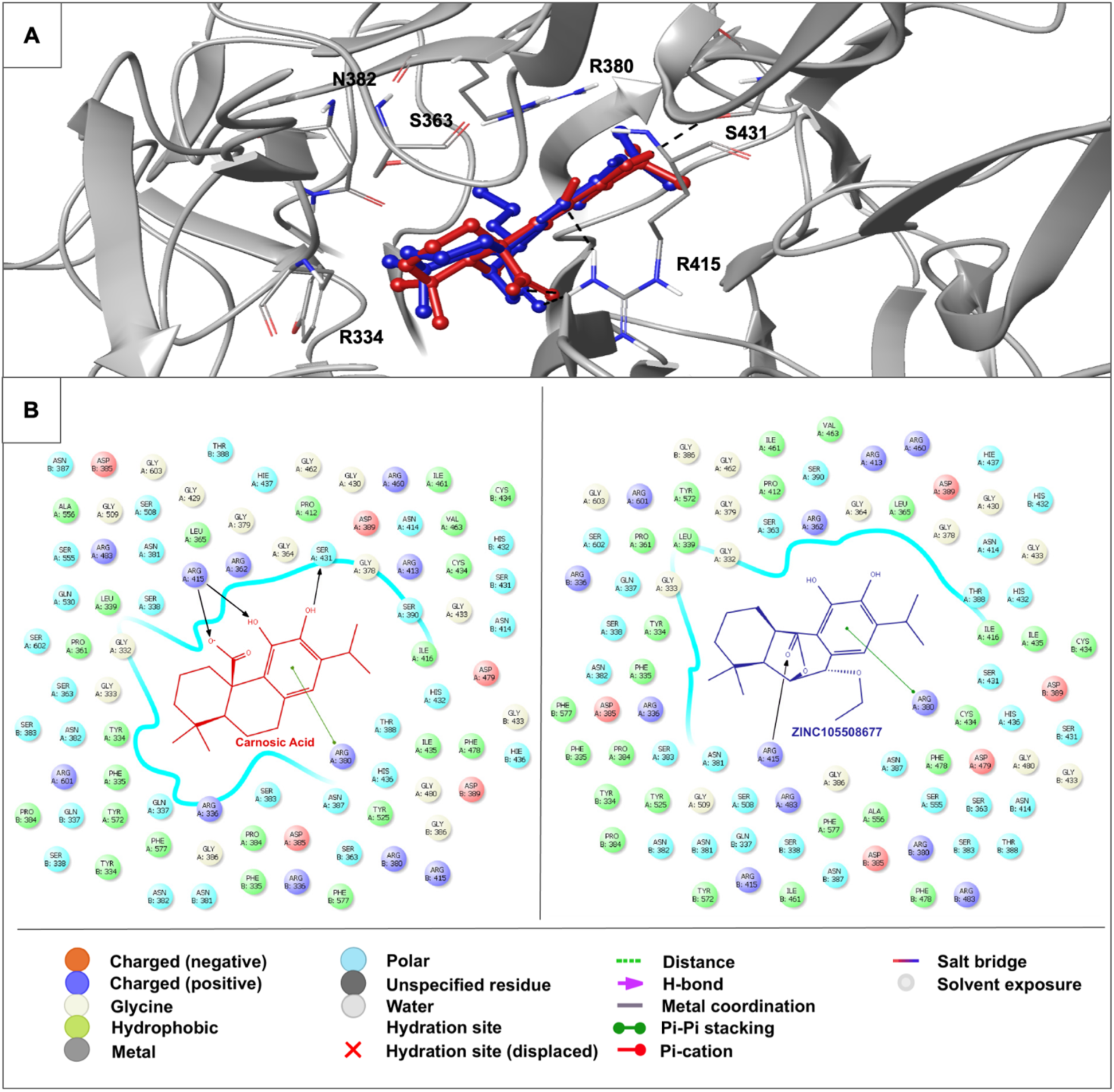
Docking results of carnosic acid and ZINC 105508677 in the binding site Keap1^Ambient_APO^ structure. (A) Docking poses of carnosic acid and ZINC 105508677 in the binding site Keap1^Ambient_APO^ structure. The light gray ribbon diagram in a metallic preset was displayed for all residues. The key residues were represented as sticks and colored by atom type: (C gray, O red, N blue). H-bond and π–π stacking interactions were outlined with dotted black and green lines, respectively. (B) Docking interactions of carnosic acid and ZINC 105508677 (colored in red and blue, respectively) in the binding site of Keap1^Ambient_APO^ structure.

### Molecular docking studies suggest CNN as the most promising drug candidate compared to L-carnosine derivatives

We obtained the most significant molecular docking results with our previously synthesized compound, CNN ***(Noguchi et al., 2019***). As CNN is an antioxidant *L*-carnosine derivative, we also performed *in silico* studies for *L*-carnosine and 100 *L*-carnosine derivatives. According to the results, CNN demonstrated the most promising results in the current study. The docking score of CNN was observed as -11.532 kcal/mol, followed by the values of the most promising *L*-carnosine derivative ZINC 450609947 (-10.529 kcal/mol) and *L*-carnosine (-9.835 kcal/mol). CNN formed strong hydrogen bonding and π cation with Arg336, Asn382, Tyr334, and Tyr572 at the ionized state. The imidazole ring and carbonyl groups greatly contributed to the high affinity of CNN and *L*-carnosine. In contrast, ZINC 450609947 presented a key hydrogen bonding through its pyrazolidinone and carboxamido moieties (**Figure 8A and 8B**).

**Figure 8.**
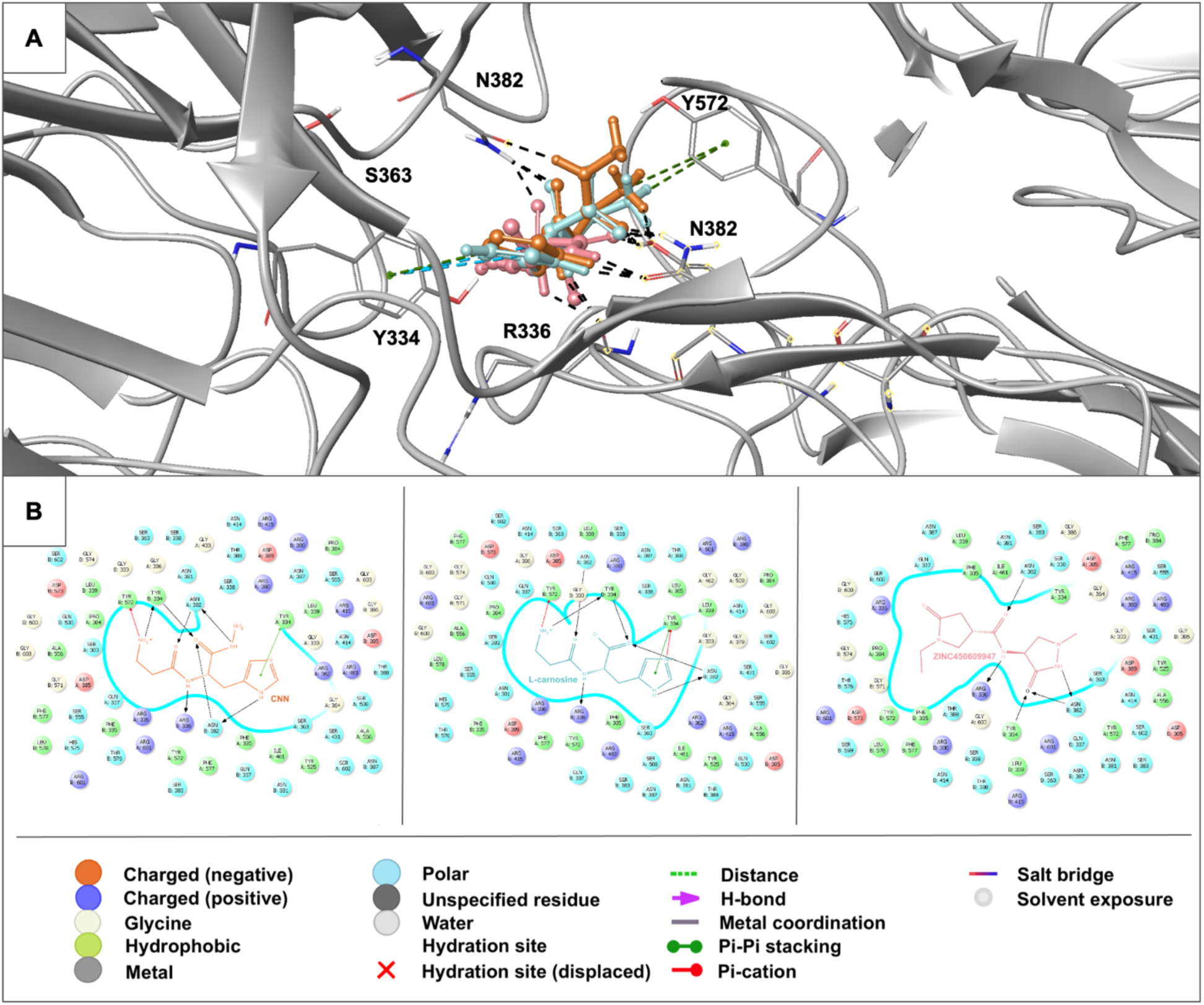
Docking results of L-carnosine, CNN and ZINC 450609947 in the binding site of Keap1^Ambient_APO^ structure. (A) Docking poses of L-carnosine, CNN and ZINC 450609947 in the binding site of Keap1^Ambient_APO^ structure. The light gray ribbon diagram in a metallic preset was displayed for all residues. The key residues were represented as sticks and colored by atom type: (C gray, O red, N blue). H-bond, π–π stacking, and π cation interactions were outlined with dotted black, green, and blue lines, respectively. (B) Docking interactions of CNN, L-carnosine, and ZINC 450609947 (colored in orange, turquoise, and pink, respectively) in the binding site of Keap1^Ambient_APO^ structure.

## Discussion

In this study, our 3Å ambient temperature structure provides details of the dimeric form Keap1 Kelch domain. Our results suggest that the Keap1 Kelch domain dimerizes, with a slight difference between chains A and B (RMSD score of 0.21 Å). Previous studies have shown the monomeric structure of the Kelch domain at cryogenic temperatures. In the Protein Data Bank, only one structure of the dimeric Kelch domain has been released, Keap1^Cryo_APO^. However, the Kelch domain dimerization has yet to be investigated. A comparative analysis of the Keap1^Ambient_APO^ and Keap1^Cryo_APO^ structures showed an RMSD score of 0.92 Å, associated with temperature-based conformational changes. Specific residues involved in dimerization, such as P384, D385, and G386, exhibited correlated fluctuations across chains.

Previous studies have shown that full-length Keap1 exists in a dimeric form, with dimerization occurring through the BTB domain, and that the distance between the Kelch domains may be important for the Keap1-Nrf2 interaction ***(Ogura et al., 2010)***. The occurrence of the dimeric Kelch domain requires further investigation to elucidate the enthalpy-entropy-based binding dynamics of the Keap1-Nrf2 interaction.

In this work, we performed GNM and GNM-TE analysis to explore the dynamic behavior of the Kelch domain structure at ambient and cryogenic temperatures, particularly for the dimerization and DMF binding behaviors. The GNM-TE method quantitatively assesses the sources and receivers of allosteric signals within a protein. This approach pinpoints the residues that serve as the most functionally significant sources of global information. These key residues exert considerable influence and function as potent effectors in the protein’s structure ***(Hacisuleyman & Erman, 2017; Altintel et al., 2022; Ersoy et al., 2023)***.

When examining the residue correlations in the dimer structures, differences were observed in the residues, possibly affecting the dimerization. On the other hand, these residues also correspond to regions where the β-factor values between the Keap1^Ambient_APO^ and Keap1 ^Cryo_APO^ differ the most, implying that these differences may arise from thermal fluctuations.

Our study revealed alterations in key residues such as Tyr334, Ser363, Val369, Gly371, Arg483, Tyr525, Glu530, Tyr572, and Phe577 when comparing the Keap1^Ambient_APO^ with Keap1^Cryo_DMF^ and Keap1^Cryo_APO^. Specifically, we observed that residues on the top side of Keap1, including Asp382, Tyr334, Phe577, Tyr572, Glu530, Tyr525, and Arg483, and those on the bottom side, such as Val369 and Gly371, exhibited significant conformational shifts. These variations were not only observed in the DMF binding residues but also in those involved in Nrf2 interaction, including Arg483, Ser508, Ser555, Arg415, Ser602, Ser363, Arg380, and Asn382, when comparing the Keap1^Ambient_APO^ structure with Keap1^Cryo_ETGE^ and Keap1^Cryo_DLG^. The temperature-based conformational shifts in these residues may influence the binding dynamics and stability of Nrf2.

Additionally, the GNM-TE analysis displays that the allosteric potential of the DMF binding residues was better revealed on the Keap1^Ambient_APO^ structure than the Keap1^Cryo_DMF^. In this analysis, within the subset of modes where DMF binding residues are identified as entropy sources, Ser390, Val467, Asn469, and Gly605 were also recognized as significant effector residues. These residues are proposed to have potential allosteric importance in DMF binding behavior.

Most compounds effective on the Nrf2 pathway possess electrophilic characteristics. These compounds bind to Keap1 and enable Nrf2 to translocate from the cytoplasm into the nucleus. They can be exemplified as follows: fumarates acting as Michael acceptors with an enone moiety or catechol-type compounds such as carnosic acid, which can acquire electrophilic potential by diverse mechanisms ***(Satoh et al., 2008; Lu et al., 2016)***. These electrophiles are irreversible indirect inhibitors of the Keap1/Nrf2 complex. This unspecific binding might cause less modulatory effects on the target and several adverse effects. Some potent direct non-covalent inhibitors of the Keap1/Nrf2 interactions were discovered based on the fact that the Kelch binding pocket is quite polar and basic and capable of interacting with carboxylic acid residues in both the ETGE and DLG motifs. Until now, various compounds carrying tetrahydroisoquinoline, 1,4-diaminonaphthalene, pyrazole, thiazole, indole, and triazole cores have been determined as potent Keap1-Nrf2 inhibitors. The carboxylic acid moiety was commonly introduced to the main structure for high efficacy with the basic Kelch binding pocket in these small molecules. However, the poor blood−brain barrier (BBB) permeability and propensity for conjugation reactions leading to reactive and toxic metabolites of carboxylic acids have shifted this approach into the replacement of the acidic groups with carboxamides and other bioisosteres ***(Pallasen et al., 2018; Ontoria et al., 2020; Sun et al., 2022; Barreca et al., 2023)***.

*L*-carnosine is an endogenous antioxidant and radical scavenger dipeptide composed of β-alanine and *L*-histidine. It possesses metal chelating, anti-inflammatory, and neuroprotective properties, which makes it suitable for investigation in disorders such as metabolic, cardiovascular, and neurodegenerative diseases. The underlying mechanism of anti-inflammatory, antioxidant, antiglycation, and anti-carbonyl effects of *L*-carnosine was mainly attributed to the activation of the Nrf2 pathway. However, the exact connection between carnosine and Nrf2 expression is still unclear ***(Aldini et al., 2021; Zhou et al., 2021; Caruso et al., 2022;2023)***. The effects of *L*-carnosine on the Nrf2 pathway could be relevant to its interactions with Keap1. In previous work, our research group synthesized the hybrid of *L*-carnosine and *L*-histidyl hydrazide (CNN). We previously reported that CNN displayed significant 4-hydroxy-trans-2-nonenal (4-HNE) scavenging activity and diminished delayed neuronal death in the hippocampus ***(Noguchi et al., 2019)***.

We also performed molecular docking studies for different fumarate derivatives, *L*-carnosine, CNN, and other *L*-carnosine derivatives, and carnosic acid and its derivatives in the Kelch domain of Keap1. CNN achieved the most promising docking results in the Keap1 Kelch domain. It formed strong interactions with key residues via both *L*-carnosine and *L*-histidyl hydrazide parts, which also interprets the high binding potential of *L*-carnosine. Fumarate derivatives bind to the Kelch domain more effectively than well-known fumarates such as DMF, ITA, MMF, MEF, and FUM. In particular, the outcome of the molecular docking of ZINC 12433145 in the Kelch domain indicated that the conversion of one of the alkyl esters to a lactone (cyclic ester) could increase the affinity of fumarate derivatives. Although ZINC 12433145 and DMF revealed similar binding profile patterns, the distances of hydrogen bonding between ZINC 12433145 and Tyr334 were found to be greater than that of DMF, which increased occupation of the whole ligand in the Kelch domain, leading to higher affinity. On the other hand, carnosic acid and its derivatives showed less binding efficacy than all tested compounds despite their prominent carboxylic acid and/or ester groups, respectively.

In conclusion, our study provides a comprehensive analysis of the dimeric Keap1 Kelch domain, emphasizing the differences in structural dynamics between ambient and cryogenic temperatures and their implications for Keap1-Nrf2 interactions. The observed temperature-based conformational shifts, particularly in key residues involved in dimerization as well as Nrf2 and DMF binding, suggest potential inferences for the binding dynamics and stability of the Keap1-Nrf2 complex. The GNM and GNM-TE analyses uncover crucial residues contributing to the allosteric effects and binding behaviors of various compounds. Remarkably, CNN, a hybrid of L-carnosine and L-histidyl hydrazide, demonstrates higher binding affinity and potential as an effective inhibitor compared to fumarate and carnosic acid derivatives. Our findings underline the importance of further exploring the enthalpy-entropy-based dynamics of Keap1 to understand its interaction with Nrf2 better and to guide the development of more effective therapeutic agents targeting the Keap1-Nrf2 pathway.

## Acknowledgments

The authors gratefully acknowledge the use of the services and the Turkish Light Source (*Turkish DeLight*) X-ray facility at Sağlık Bilimleri Üniversitesi Deneysel Tıp Araştırma ve Uygulama Merkezi (SBU-DETUAM). The authors also gratefully acknowledge the use of the services and facilities at KoçUniversity Isbank Research Centre for Infectious Diseases (KUIS-CID). The authors declare no conflicts of interest. This publication has benefited from the 2232 International Fellowship for Outstanding Researchers Program and the 1001 Scientific and Technological Research Projects Funding Program of the TÜBİTAK (Project Nos. 118C270, 120Z520, and 119F392). However, the entire responsibility of the publication belongs to the authors of the publication. The financial support received from TÜBİTAK does not mean that the content of the publication is approved in a scientific sense by TÜBİTAK.

