## Supplementary Figures for "Insights into Therapeutic Discovery Through the Kelch Domain Structure of Keap1 at Ambient Temperature"

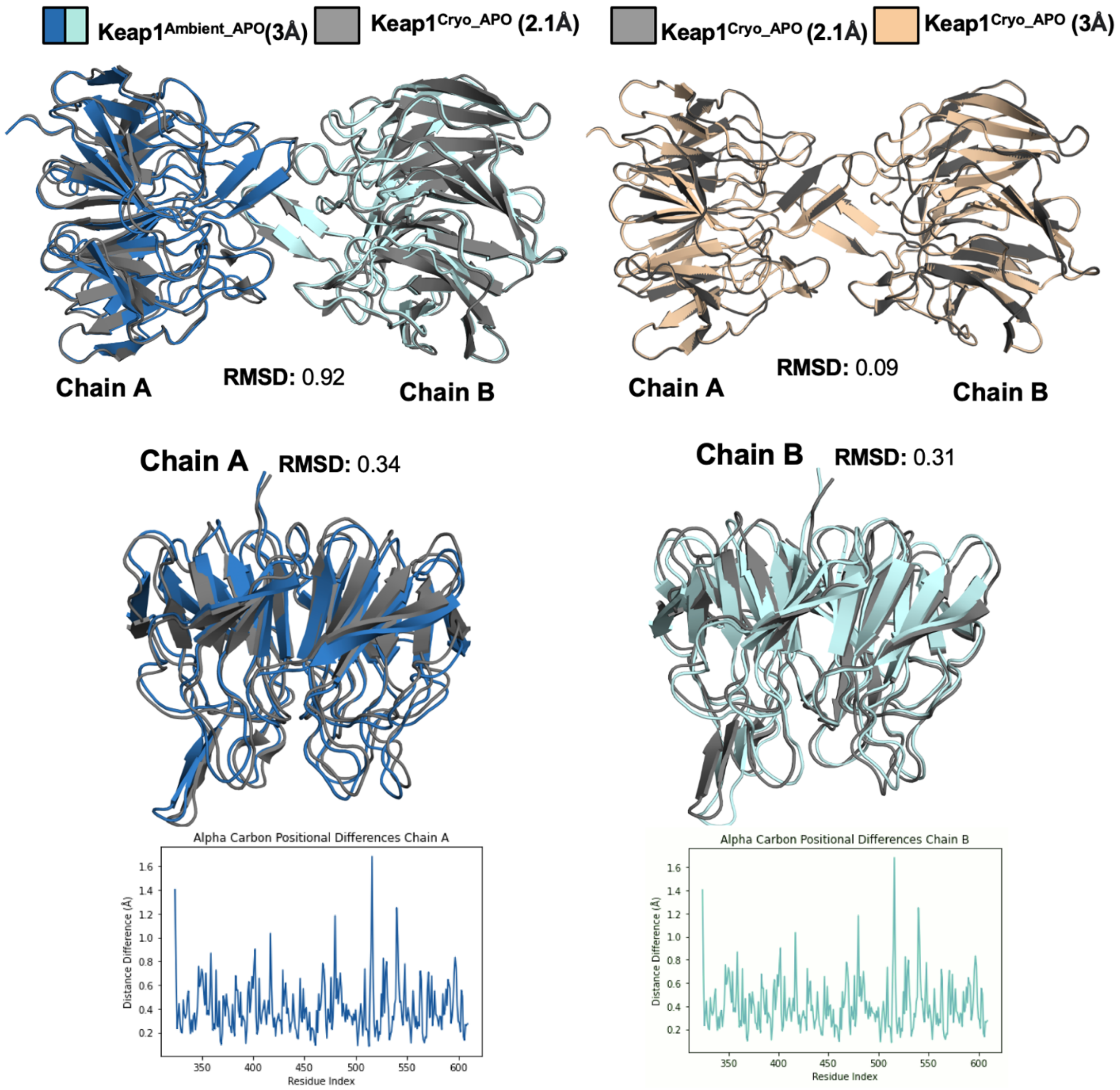


***Supplementary Figure 1.*** *Superposition of 2.16 Å* Keap1$^{Cryo_APO}$ *(colored in gray50) and 3 Å* Keap1$^{Ambient_APO}$ *(colored in sky blue and pale cyan according to monomers), with an RMSD score of 0.92 Å. Superposition of 2.16 Å* Keap1$^{Cryo_APO}$ *and 3 Å* Keap1$^{Cryo_APO}$ *(colored in wheat), with an RMSD score of 0.09 Å. Alignment of 2.16 Å* Keap1$^{Cryo_APO}$ *and* Keap1$^{Ambient_APO}$ *Chain A with an RMSD score of 0.34 Å, demonstrated by a pairwise distance plot. Alignment of 2.16 Å* Keap1$^{Cryo_APO}$ *and* Keap1$^{Ambient_APO}$ *Chain B with an RMSD score of 0.31 Å, demonstrated by a pairwise distance plot.*

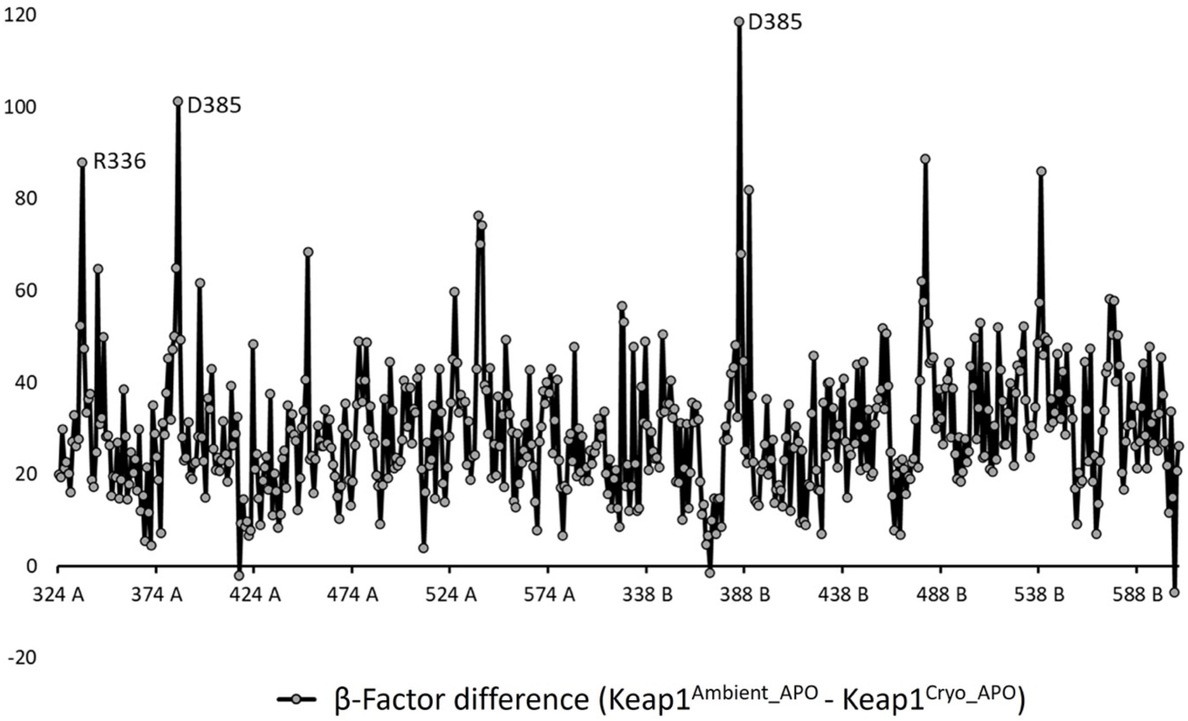


**Supplementary Figure 2.** *β-factor differences between the two dimeric structures obtained at ambient* (Keap1$^{Ambient_APO}$) and *cryogenic* (Keap1$^{Cryo_APO}$) *temperatures.*

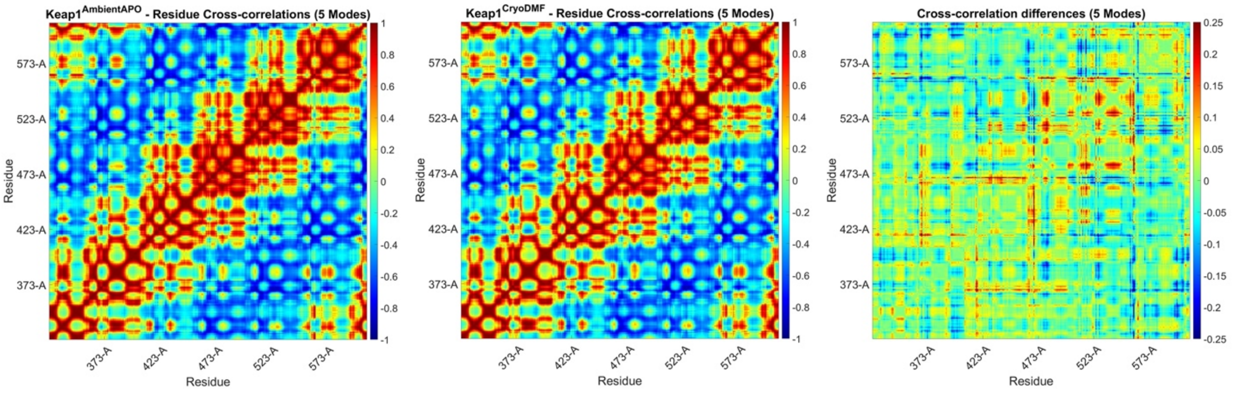


**Supplementary Figure 3.** *GNM residue cross-correlations for five slowest GNM modes of the monomer Keap1 Kelch domain structures* Keap1$^{Ambient_APO}$ (Chain A) *and the* Keap1$^{Cryo_DMF}$ (Monomer)*, together with the differences of the residue cross correlations. Only two residues (N469 & I559) show slight differences in correlations.*

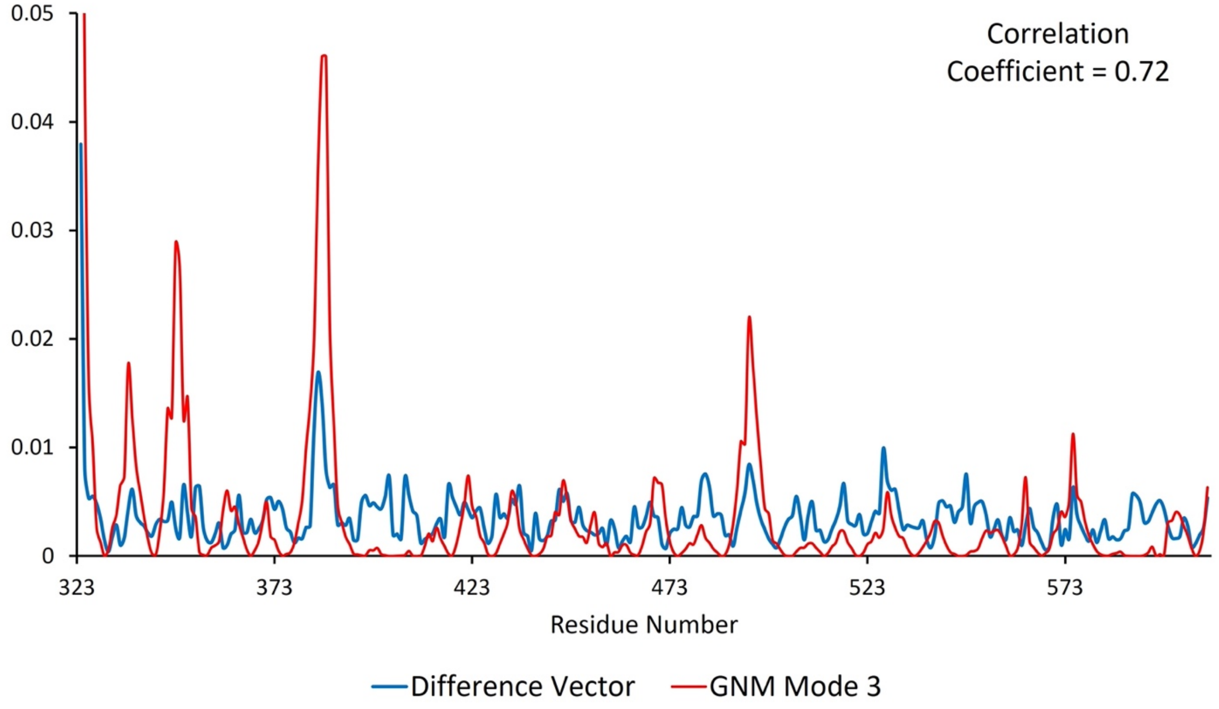


| | Mode-1 | Mode-2 | Mode-3 | Mode-4 | Mode-5 |
|---|---|---|---|---|---|
| **Correlation Coefficient** | 0.36 | 0.03 | **0.72** | 0.49 | 0.03 |

**Supplementary Figure 4**. *Mean squared residue fluctuations of the third slowest mode of the kelch domain* of Keap1$^{Cryo_DMF}$ *and the difference vector between* Keap1$^{Cryo_DMF}$ *and* Keap1$^{Ambient_APO}$ (Chain A). Together with the correlation coefficient between each of the five slowest GNM modes and the difference vector.

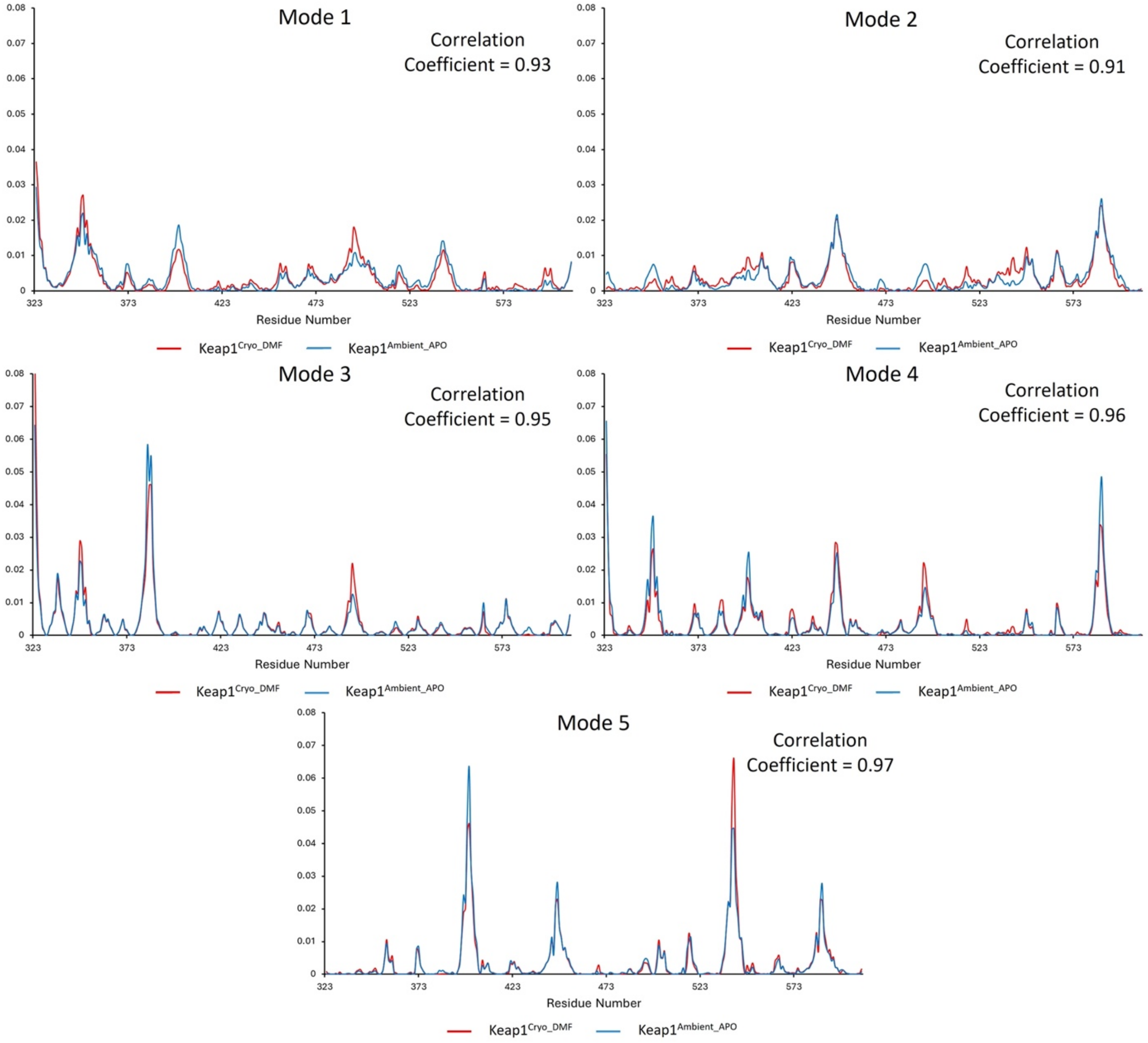


**Supplementary Figure 5.** *Mean squared residue fluctuations of the five slowest GNM modes of the Keap1 Kelch domain structures obtained at cryogenic* ($\text{Keap1}^{\text{Cryo\_DMF}}$) *and ambient* ($\text{Keap1}^{\text{Ambient\_APO}}$ Chain A) *temperature. Correlation coefficients between slow modes are displayed at the top right corner of the graphs.*

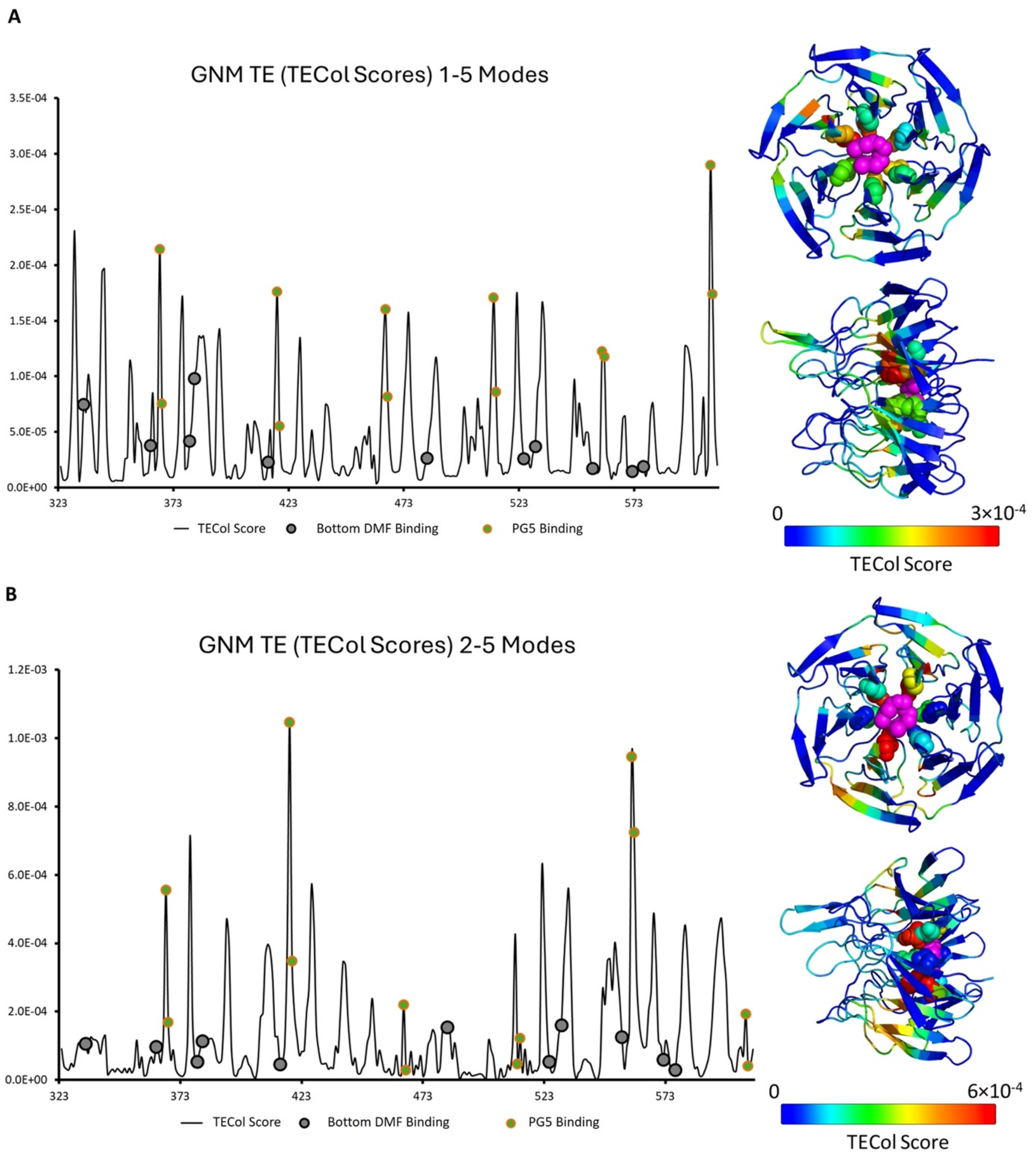


**Supplementary Figure 6.** *The TECol score results with a subset of slow modes consisting of GNM modes 1 to 5 (A) and 2 to 5 (B) for the monomer Keap1 Kelch domain obtained at ambient temperature* (Keap1$^{Ambient_APO}$ chain A). *DMF binding and PG5 binding residues are marked on the graphs. 3D representation of the TECol score results are given from two angles. Residues are*

*colored according to TECol score with respect to the rainbow spectrum. PG5 binding residues are shown as spheres and PG5 is shown as magenta sphere.*

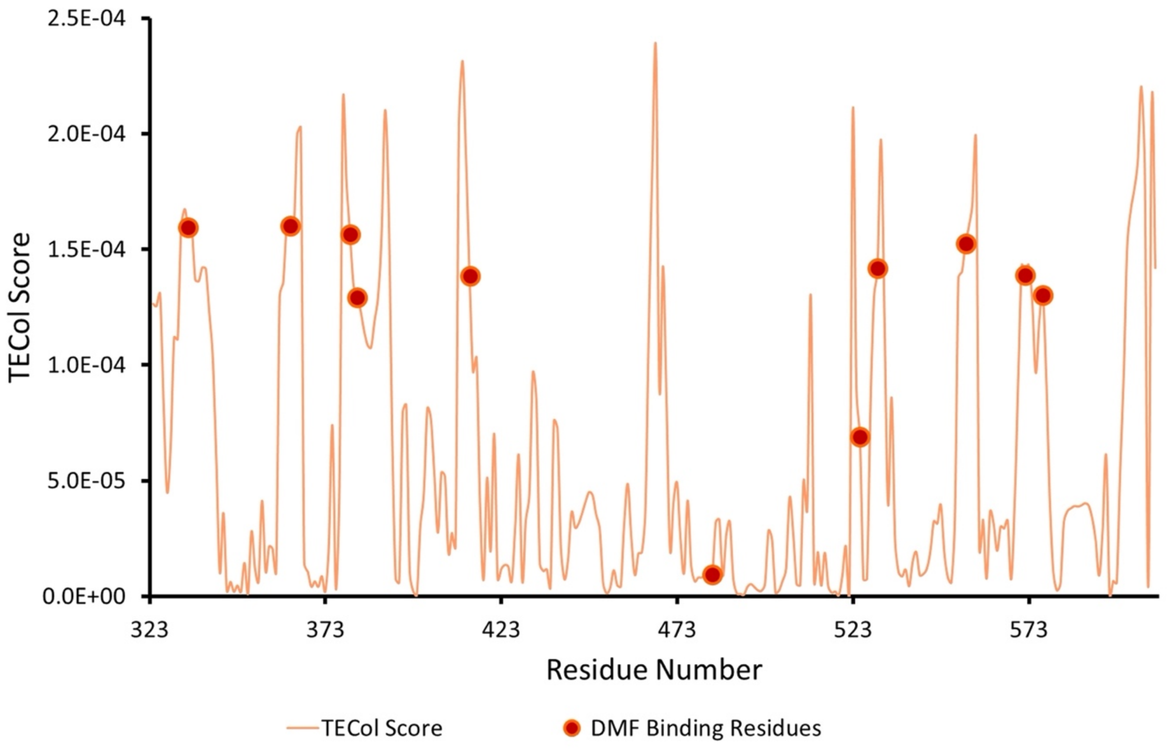


**Supplementary Figure 7.** *The TECol score results with a subset of slow modes consisting of GNM modes 3 to 5 for the monomer Keap1 Kelch domain obtained at cryogenic temperature* (Keap1$^{Cryo_DMF}$).

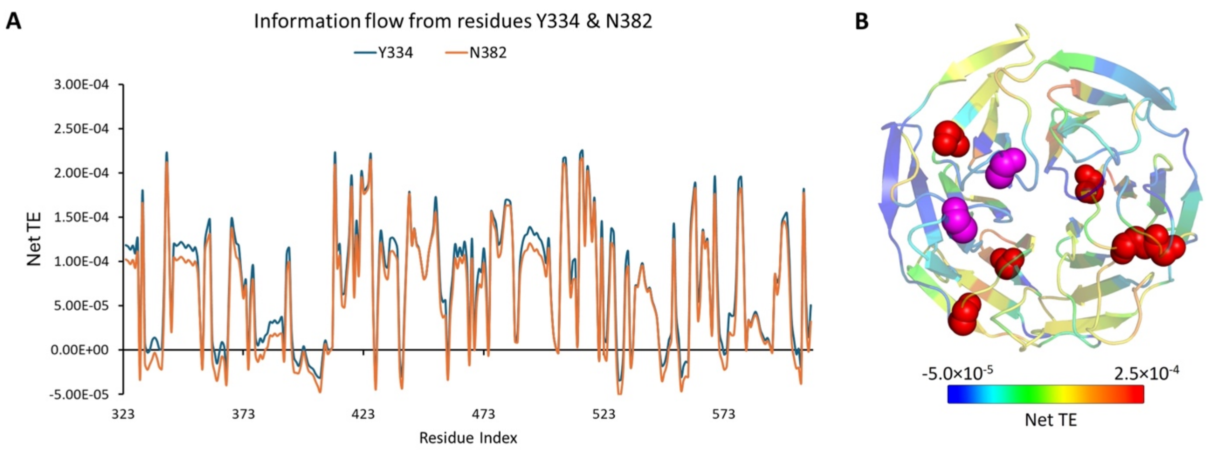


**Supplementary Figure 8.** *(A) 2D graph of the information flow from residues Y334 and N382 obtained from GNM-TE analysis with the subset of slow modes consisting of GNM modes 3 to 5 for the monomer Keap1 Kelch domain obtained at ambient temperature* (Keap1$^{Ambient_APO}$ chain A). *(B) 3D representation of the information flow from residue Y334 where the residues are colored according to Net TE values with respect to the rainbow spectrum. The residues that receive the highest information (Y342, V411, Y426, I506, R507 and V514) are represented as spheres and residues Y334 and N382 and represented as magenta spheres.*
